# A Convolutional Autoencoder-based Explainable Clustering Approach for Resting-State EEG Analysis

**DOI:** 10.1101/2023.01.04.522805

**Authors:** Charles A. Ellis, Robyn L. Miller, Vince D. Calhoun

## Abstract

Machine learning methods have frequently been applied to electroencephalography (EEG) data. However, while supervised EEG classification is well-developed, relatively few studies have clustered EEG, which is problematic given the potential for clustering EEG to identify novel subtypes or patterns of dynamics that could improve our understanding of neuropsychiatric disorders. There are established methods for clustering EEG using manually extracted features that reduce the richness of the feature space for clustering, but only a couple studies have sought to use deep learning-based approaches with automated feature learning to cluster EEG. Those studies involve separately training an autoencoder and then performing clustering on the extracted features, and the separation of those steps can lead to poor quality clustering. In this study, we propose an explainable convolutional autoencoder-based approach that combines model training with clustering to yield high quality clusters. We apply the approach within the context of schizophrenia (SZ), identifying 8 EEG states characterized by varying levels of δ activity. We also find that individuals who spend more time outside of the dominant state tend to have increased negative symptom severity. Our approach represents a significant step forward for clustering resting-state EEG data and has the potential to lead to novel findings across a variety of neurological and neuropsychological disorders in future years.

## I. Introduction

The application of machine learning to resting-state electroencephalograms (EEG) has grown increasingly common. The domain of EEG-based supervised classification is highly developed [1][2]. In contrast, the domain of unsupervised EEG clustering is still largely in its infancy [3][4]. This is partly due to the limited number of clustering approaches for lengthy EEG time-series and is problematic because clustering EEG offers key opportunities. Namely, clustering can identify novel disorder-related dynamics or subtypes, as has been shown in EEG [3] and functional magnetic resonance imaging (fMRI) [5][6] analysis. Moreover, most EEG clustering methods do not give insight into the features important to their clusters (i.e., they are not explainable). In this study, we present a novel explainable autoencoder-based clustering approach that can uncover disorder-related dynamics in resting-state EEG. We demonstrate its viability within the context of schizophrenia (SZ).

EEG clustering involves (1) traditional clustering with engineered features or (2) deep learning with raw data and automated feature extraction. Feature engineering methods are used in most studies [3], [4]. However, extracted feature limit the information available for learning. Deep learning, in contrast, could enable a more thorough exploration of EEG through automated feature extraction. This is particularly important when trying to uncover novel aspects of neurological and neuropsychiatric disorders. EEG deep learning-based clustering mainly uses autoencoders [7][8]. Autoencoders can perform automated feature extraction and have been used in EEG classification [9][10]. When used for clustering, autoencoders extract a condensed representation of the original data that is then clustered via traditional algorithms [7][8]. This approach has an important shortcoming. Specifically, the feature extraction and clustering are separate processes. This can result in the extraction of features that are linearly inseparable and yield low quality clusters. Jointly optimizing feature extraction and clustering could increase cluster quality [11], [12].

Additionally, if clustering methods are to be applied for insight into EEG data, it would be helpful if they could identify the features most important to the clustering. To our knowledge, no EEG clustering methods have integrated explainability approaches. Even among clustering algorithms that use engineered features, explainability is limited, and the field is in its infancy [13]. Moreover, to our knowledge, no methods have been developed to explain deep learning-based clustering.

In this study, we present a novel explainable convolutional autoencoder clustering approach for resting-state EEG. It combines both autoencoder training and clustering into a single process that jointly optimizes extracted cluster quality and reconstruction loss. It also gives insight into the spectral features critical to the clustering. We demonstrate the viability of the approach within the context of SZ. We identify multiple neurological states in the resting-state EEG before extracting features related to the dynamics of the states and analyzing their relationship with SZ symptom severity. Our study provides a novel approach for the domain of resting-state EEG analysis and represents a significant step forward for EEG clustering.

## II. Methods

### A. Description of Data and Preprocessing

We used a resting-state scalp EEG dataset with 47 individuals with SZ (SZs) [14]. The dataset has been used for multiple classification studies [1]. All study participants gave written informed consent, and data collection was approved by the Hartford Hospital Internal Review Board. The standard 10-20 format of 64 electrodes was used during collection. Like previous studies [1], we only used 19 electrodes. Data was recorded at 1000 Hertz (Hz) for 5 minutes. We downsampled the data to 200 Hz and used subject-specific min-max normalization on a per-channel basis. We segmented the data into 5-second segments using a sliding window approach with a 2.5-second overlap. We then channel-level z-scored each sample. Lastly, we used data augmentation to increase the number of samples. We duplicated the dataset twice and added Gaussian noise with a mean of 0 and a standard deviation of 0.3 to each duplicate. This effectively tripled the dataset size. The dataset also contained information on symptom severity from the Positive and Negative Syndrome Scale (PANSS) [5] with scores for positive, negative, and general symptoms.

### A. Convolutional Autoencoder-based Clustering Approach for State Identification

Figure 1 shows the autoencoder architecture. We initially tuned the hyperparameters by optimizing the mean absolute error (MAE) of the autoencoder across all samples. After obtaining a satisfactory reconstruction loss, we created a multi-objective, composite loss function (Equation 1) and updated our training approach. Similar approaches were presented in [11][12]. Our training approach had several steps. (1) The autoencoder weights were initialized. (2) Cluster centroids (***C***) were randomly initialized, where centroids were in the space of extracted features. (3) The model weights were updated with minibatch gradient descent (batch size = 128) using the composite loss. (4) After an epoch, k-means clustering was performed on the extracted features, and the cluster centroids were updated. (5) Steps 3 and 4 were repeated for 100 epochs.

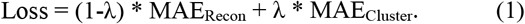

**Figure 1.**
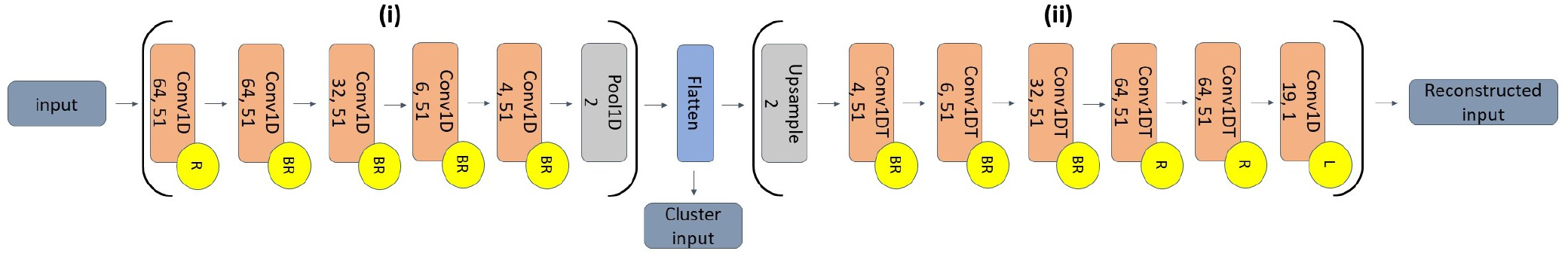
Convolutional Autoencoder Architecture. The architecture has two segments - (i) an encoder and (ii) a decoder – that are separated by a flatten layer. Conv1D and Conv1DT layers are 1D convolutional and 1D convolutional transpose layers, respectively. The first and second numbers below the layer type are the number of filters and filter size. Pool1D is a max pooling layer with a pooling size of 2. Upsample is an upsampling layer with a factor of 2. “R”, “L”, and “B” in circles to the bottom right of layers indicate that those layers have ReLU and linear activation functions and batch normalization layers. Layers did not use any padding, and all Conv1D and Conv1DT layers used He Normal initialization with strides of 1.

***MAE_Recon_*** is the MAE of the reconstructed data relative to the original data. ***MAE_Cluster_*** is the MAE between the features extracted for samples at the flatten layer and its corresponding cluster centroid. We used a grid search varying the number of clusters from 2 to 10 and varying the *λ* parameter from 0.1 to 0.9 in 0.1-unit increments. For each parameter set, we selected the model from the epoch with the lowest loss. We then estimated cluster quality using the Calinksi-Harabasz (CH) and Davies-Bouldin (DB) scores. We balanced the need for both high cluster quality and reconstruction when selecting an optimal parameter set. We ranked the parameters based upon the cluster quality metrics and reconstruction error. We averaged the cluster quality rankings and then averaged the combined cluster quality ranking with the reconstruction error ranking. To visualize the extracted features, we applied Principal Component Analysis (PCA).

### C. Spectral Explainability for Insight into Identified States

After identifying each of the EEG states, we wanted to identify the important features for each state. To this end, we first performed a simple visualization by converting the samples assigned to each state to the frequency domain via a Fast Fourier Transform (FFT), feature-wise z-scoring, and averaging for comparison. We then applied a novel spectral explainability approach for insight into the key frequency bands differentiating each state. Our approach combines a version of EasyPEASI [15] with a metric from [13]. EasyPEASI [16] is typically used to explain supervised raw EEG classifiers and involves perturbing EEG frequency bands and examining their effect on classifier performance. Our approach involves several steps. (1) Samples are assigned to a cluster. (2) Samples are converted to the frequency domain via an FFT. (3) Coefficients from a frequency band are permuted across samples. (4) Perturbed samples are converted back to the time domain via an inverse FFT. (5) The perturbed samples are reassigned to the existing clusters. (6) The percentage of samples that switch clusters is calculated, and (7) steps 2 through 6 are repeated for each frequency band. The process is repeated multiple times to obtain a distribution of the effects of perturbing each frequency band. We perturbed each frequency band 100 times, and we used the frequency bands: δ (0 – 4 Hz), θ (4 – 8 Hz), α (8 – 12 Hz), β (12 – 25 Hz), γ_lower_ (25 – 55 Hz), and γ_upper_ (65 – 100 Hz).

### D. Statistical Analysis to Identify Relationships between Identified States and Symptoms

We wanted to relate the identified states to SZ symptom severity. To this end, we used a metric frequently used in fMRI analysis [5], [17]. Namely, we calculated the occupancy rate (OCR, i.e., the percent of samples belonging to the dominant state) for each participant. We then used ordinary least squares regression, controlling for age and sex, to estimate the relationship between symptom severity and OCR.

## III. Results and Discussion

In this section, we describe and discuss the results of our explainable clustering analysis. We also discuss limitations of our approach and future research directions.

### A. Identifying EEG States

Eight clusters at *λ*=0.3 was the best parameter set. Figure 2 shows the clustering results. The composite loss decreased as *λ* increased, indicating that reducing ***MAE_Recon_*** was harder than obtaining condensed clusters. The ***MAE_Recon_*** increased with *λ*. Cluster quality tended to be highest when *λ* was away from the extremes of 0.1 and 0.9. The ranking prioritized mid-*λ* values. As such, *λ* being too high likely caused dense clusters to form before effective representations could be extracted. Importantly, lower *λ* values that simulate preexisting autoencoder clustering often had lower cluster quality, which supports the viability of our approach for improving cluster quality.

**Figure 2.**
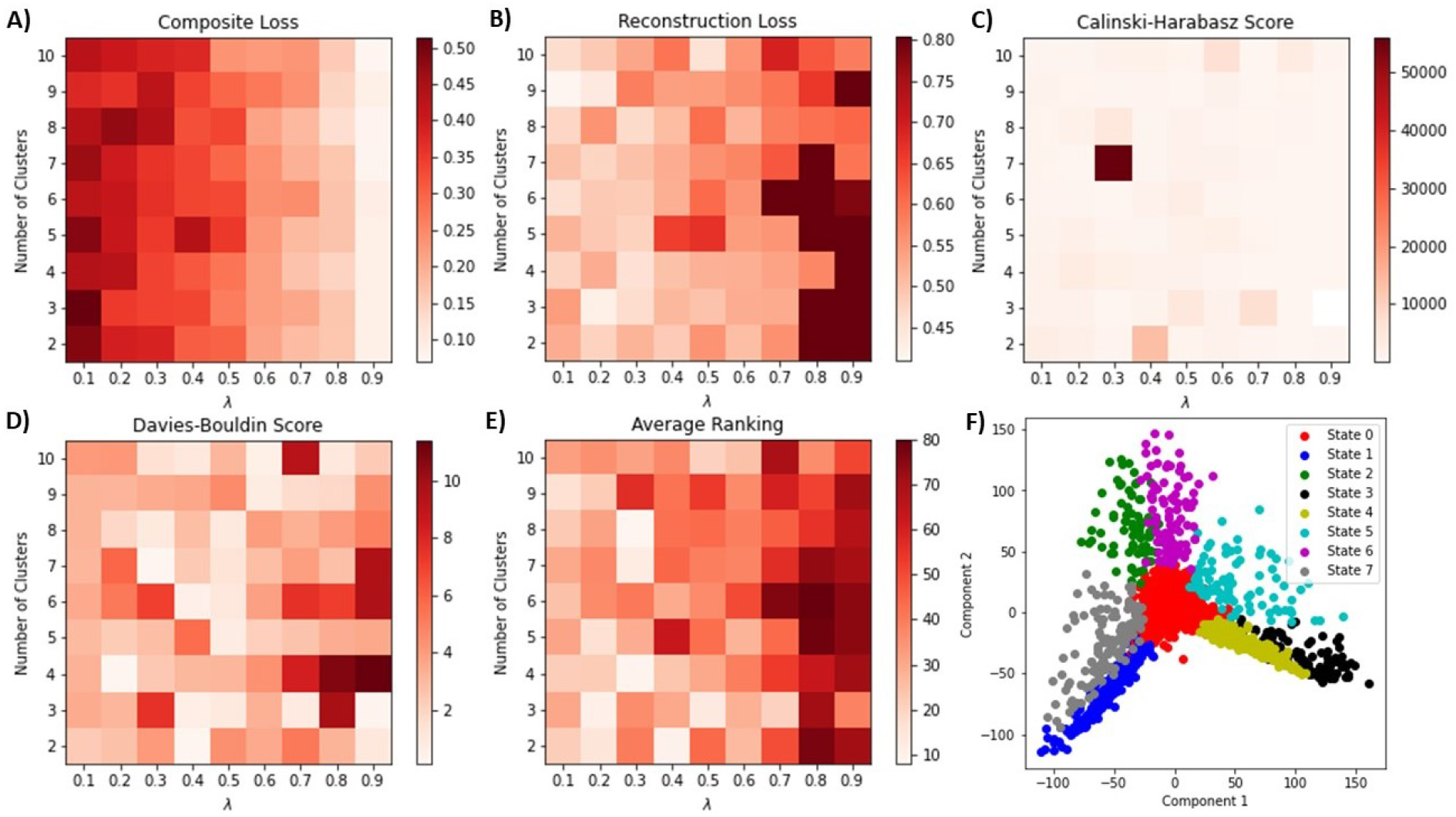
Cluster Quality and PCA Results. Panels A and B show the autoencoder composite loss and reconstruction loss. Panels C, D, and E show the CH score, DB score, and average ranking, respectively. Panel F shows two principal components of the extracted features for the unaugmented data. In panels A through E, the x and y-axes show the lambda values and number of clusters, respectively. A color bar to the right of each panel shows the values associated with their color maps. In panel F, samples in each state are each shown in a different color. Higher CH and lower DB scores are better.

### B. Characterizing EEG States

Figure 2F shows the principal components of the features extracted at the bottleneck layer and indicates that individuals mainly spent time in a core state (State 0) and periodically migrated to surrounding states (States 1 through 7). Figure 3 shows the average z-scored spectral power across channels in each identified state. State 0 had moderate spectral power. Other states tended to have relatively high or low power. Figure 4 shows the results of our spectral importance analysis. Interestingly, only the δ band seemed to have a strong effect upon the clustering, though θ also had some effect. Previous studies have found relationships between δ and SZ [1].

**Figure 3.**
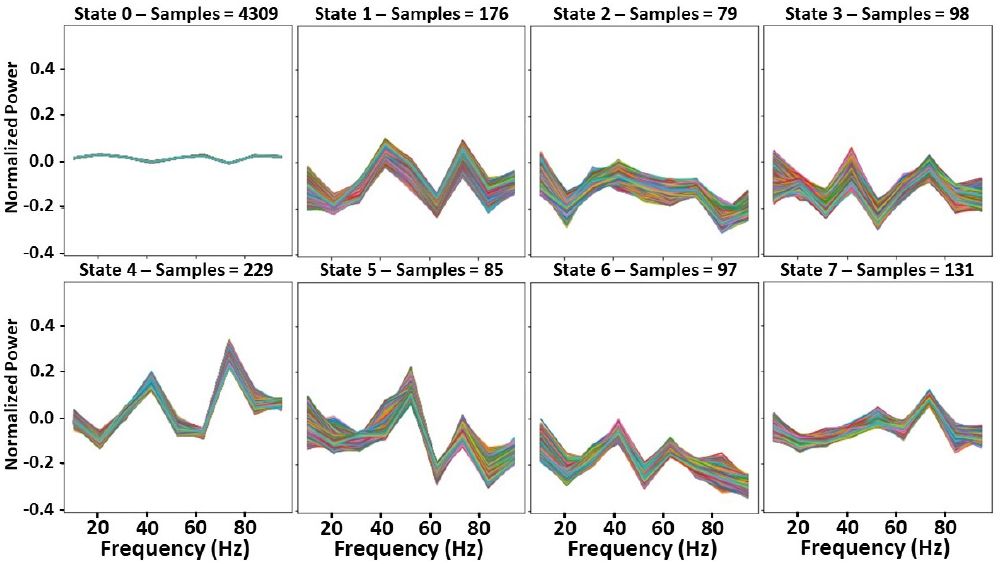
Spectral Power for Each State. The x-axis of each panel indicates the frequency in Hertz, and the y-axis indicates normalized power. The state and corresponding number of samples in the title of each panel.

**Figure 4.**
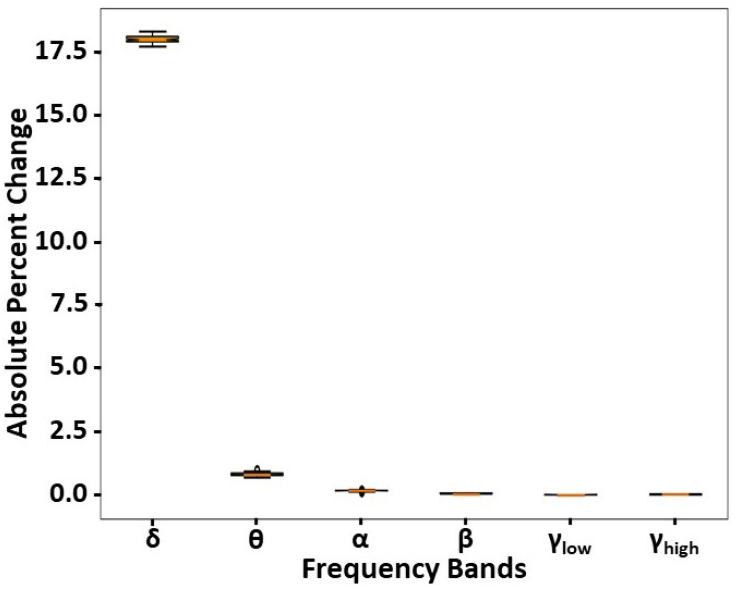
Spectral Importance Estimates. The x-axis indicates each of the perturbed frequency bands, and the y-axis indicates the absolute percent change in cluster assignments following perturbation.

### C. Identifying Relationships Between States and Symptoms

Based on the ordinary least squares regression controlling for age and gender, we found a significant inverse relationship between negative symptom severity and state 0 OCR (p < 0.01) but did not identify significant relationships between positive and general symptoms and state 0 OCR. Our findings indicate that SZs who spent more time in states with relatively high or low spectral power tended to have worse negative symptom severity.

### D. Limitations and Next Steps

Our characterization of the identified states primarily focused on identifying their spectral features. It would be interesting to examine the importance of individual channels [18] or to examine key waveforms of relevance to the clustering [19][20]. The δ importance that we identified could be explained by how the expected amplitude of EEG signal tends to decrease as its frequency increases. As such, higher amplitude, and thus lower frequency, activity would be penalized more than higher frequency activity by reconstruction loss. This will likely be problematic for any autoencoder-based clustering study and could be addressed by reducing reconstruction loss.

## IV. Conclusion

Clustering resting-state EEG could help identify novel neuropsychiatric disorder subtypes or dynamics. Some studies have used convolutional autoencoders to cluster EEG. However, their disjointed model training and clustering process leads to low quality clustering. In this study, we present a novel explainable clustering approach that combines model training and clustering to obtain high quality clusters. We apply our method to SZ, identifying a dominant state of moderate δ power in which SZs with higher negative symptom severity spend less time. Our approach is a significant step forward for EEG clustering and has the potential to be applied for novel insights into a variety of neuropsychiatric and neurological disorders.

## References

[1] C. A. Ellis, A. Sattiraju, R. Miller, and V. Calhoun, “Examining Effects of Schizophrenia on EEG with Explainable Deep Learning Models,” in 2022 IEEE 22nd International Conference on Bioinformatics and Bioengineering (BIBE), 2022, pp. 301–304, doi: 10.1109/BIBE55377.2022.00068.

[2] C. A. Ellis, A. Sattiraju, R. Miller, and V. Calhoun, “Examining Reproducibility of EEG Schizophrenia Biomarkers Across Explainable Machine Learning Models,” in 2022 IEEE 22nd International Conference on Bioinformatics and Bioengineering (BIBE), 2022, pp. 305–308, doi: 10.1109/BIBE55377.2022.00069.

[3] N. Gurudath and H. Bryan Riley, “Drowsy driving detection by EEG analysis using Wavelet Transform and K-means clustering,” Procedia Comput. Sci., vol. 34, pp. 400–409, 2014, doi: 10.1016/j.procs.2014.07.045.

[4] R. Lin, R.-G. Lee, C.-L. Tseng, Y.-F. Wu, and J.-A. Jiang, “Multi-channel EEG recording system and study of EEG clustering method,” Biomed. Eng. Appl. Basis Commun., vol. 18, no. 6, pp. 276–283, 2006.

[5] M. S. E. Sendi et al., “Aberrant Dynamic Functional Connectivity of Default Mode Network in Schizophrenia and Links to Symptom Severity,” Front. Neural Circuits, vol. 15, no. March, pp. 1–14, 2021, doi: 10.3389/fncir.2021.649417.

[6] C. A.. Ellis, R. L.. Miller, and V. D.. Calhoun, “Identifying Neuropsychiatric Disorder Subtypes and Subtype-Dependent Variation in Diagnostic Deep Learning Classifier Performance,” bioRxiv, pp. 2–5, 2022.

[7] K. S. Prabhudesai, L. M. Collins, and B. O. Mainsah, “Automated feature learning using deep convolutional auto-encoder neural network for clustering electroencephalograms into sleep stages,” Int. IEEE/EMBS Conf. Neural Eng. NER, vol. 2019-March, pp. 937–940, 2019, doi: 10.1109/NER.2019.8716996.

[8] H. Daoud and M. Bayoumi, “Deep Learning Approach for Epileptic Focus Localization,” IEEE Trans. Biomed. Circuits Syst., vol. 14, no. 2, pp. 209–220, 2019, doi: 10.1109/TBCAS.2019.2957087.

[9] L. Fraiwan and K. Lweesy, “Neonatal sleep state identification using deep learning autoencoders,” Proc. - 2017 IEEE 13th Int. Colloq. Signal Process. its Appl. CSPA 2017, vol. 2, no. March, pp. 228–231, 2017, doi: 10.1109/CSPA.2017.8064956.

[10] J. Liu et al., “EEG-Based Emotion Classification Using a Deep Neural Network and Sparse Autoencoder,” Front. Syst. Neurosci., vol. 14, no. September, pp. 1–14, 2020, doi: 10.3389/fnsys.2020.00043.

[11] C. Song, F. Liu, Y. Huang, L. Wang, and T. Tan, “Auto-encoder based data clustering,” in Lecture Notes in Computer Science (including subseries Lecture Notes in Artificial Intelligence and Lecture Notes in Bioinformatics), 2013, vol. 8258 LNCS, no. PART 1, pp. 117–124, doi: 10.1007/978-3-642-41822-8_15.

[12] H. Muhammad et al., “Unsupervised subtyping of cholangiocarcinoma using a deep clustering convolutional autoencoder,” Lect. Notes Comput. Sci. (including Subser. Lect. Notes Artif. Intell. Lect. Notes Bioinformatics), vol. 11764 LNCS, pp. 604–612, 2019, doi: 10.1007/978-3-030-32239-7_67.

[13] C. A. Ellis, M. S. E. Sendi, E. P. T. Geenjaar, S. M. Plis, R. L. Miller, and V. D. Calhoun, “Algorithm-Agnostic Explainability for Unsupervised Clustering,” pp. 1–22, 2021, [Online]. Available: http://arxiv.org/abs/2105.08053.

[14] J. Sui et al., “Combination of FMRI-SMRI-EEG data improves discrimination of schizophrenia patients by ensemble feature selection,” 2014 36th Annu. Int. Conf. IEEE Eng. Med. Biol. Soc. EMBC 2014, no. 2, pp. 3889–3892, 2014, doi: 10.1109/EMBC.2014.6944473.

[15] C. A. Ellis, R. L. Miller, and V. D. Calhoun, “A Novel Local Explainability Approach for Spectral Insight into Raw EEG-Based Deep Learning Classifiers,” in 21st IEEE International Conference on BioInformatics and BioEngineering, 2021, pp. 0–5.

[16] D. O. Nahmias and K. L. Kontson, “Easy Perturbation EEG Algorithm for Spectral Importance (easyPEASI): A Simple Method to Identify Important Spectral Features of EEG in Deep Learning Models,” in Proceedings of the 26th ACM SIGKDD International Conference on Knowledge Discovery & Data Mining, Aug. 2020, pp. 2398–2406, doi: 10.1145/3394486.3403289.

[17] C. A. Ellis, M. S. E. Sendi, R. L. Miller, and V. D. Calhoun, “An Unsupervised Feature Learning Approach for Elucidating Hidden Dynamics in rs-fMRI Functional Network Connectivity,” in 2022 44th Annual International Conference of the IEEE Engineering in Medicine & Biology Society (EMBC), 2022, pp. 4449–4452.

[18] C. A. Ellis et al., “Novel Methods for Elucidating Modality Importance in Multimodal Electrophysiology Classifiers,” bioRxiv, 2022.

[19] C. A. Ellis, R. L. Miller, and V. D. Calhoun, “A Model Visualization-based Approach for Insight into Waveforms and Spectra Learned by CNNs,” in IEEE, 2021, pp. 1–4.

[20] C. A. Ellis, R. L. Miller, and V. D. Calhoun, “A Systematic Approach for Explaining Time and Frequency Features Extracted by Convolutional Neural Networks From Raw Electroencephalography Data,” Front. Neuroinform., vol. 16, no. May, pp. 1–11, 2022, doi: 10.3389/fninf.2022.872035.

